# Purinergic P2Y1 and P2Y2/4 receptors elicit distinct Ca^2+^ signaling patterns in hepatocytes via differential feedback regulation by Protein Kinase C

**DOI:** 10.1101/2020.10.29.360784

**Authors:** Juliana C. Corrêa-Velloso, Paula J. Bartlett, Robert Brumer, Lawrence Gaspers, Henning Ulrich, Andrew P. Thomas

**Affiliations:** Department of Pharmacology, Physiology & Neuroscience, New Jersey Medical School, Rutgers, The State University of New Jersey, USA; Department of Biochemistry, Institute of Chemistry, University of São Paulo, São Paulo, Brazil

**Keywords:** purinergic receptors, liver, protein kinase C (PKC), feedback inhibition, calcium, Ca^2+^ oscillations, ADP, UTP

## Abstract

Extracellular nucleotides are key regulators of liver physiology. In primary rat hepatocytes, P2Y1 receptor (P2Y1R) activation by ADP generates cytosolic calcium ([Ca^2+^]_c_) oscillations with narrow spikes, whereas P2Y2/4R activation by UTP led to more complex broad [Ca^2+^]_c_ oscillations. Both [Ca^2+^]_c_ oscillation signatures were observed with the common agonist ATP. Inhibition of Gα_q_ signaling with YM-254890 abolished ATP-induced [Ca^2+^]_c_ oscillations, indicating that they depend on inositol 1,4,5-trisphosphate (IP_3_), and are not mediated by P2X receptors. The narrow P2Y1-linked [Ca^2+^]_c_ spikes and the broad P2Y2/4-linked [Ca^2+^]_c_ spikes are shaped by differential and complex PKC-mediated feedback mechanisms. Downregulation of PKC broadened both ADP- and UTP-induced [Ca^2+^]_c_ oscillations, with a more pronounced effect on the former. PKC downregulation also selectively elicited a more robust response to ADP stimulation, enhancing oscillatory and sustained [Ca^2+^]_c_ responses. Acute PKC modulation confirmed the importance of the negative PKC feedback regulation of P2Y1R-linked [Ca^2+^]_c_ signals; such that PKC activation decreased [Ca^2+^]_c_ oscillation frequency and PKC inhibition increased [Ca^2+^]_c_ spike width. However, both PKC activation and inhibition decreased the spike width of P2Y2/4R-induced [Ca^2+^]_c_ oscillations, suggesting that multiple opposing PKC feedback mechanisms shape P2Y2/4R responses. Significantly, plasma membrane Ca^2+^ entry was required for negative PKC feedback on P2Y1R-linked [Ca^2+^]_c_ oscillations, whereas P2Y2/4R-linked [Ca^2+^]_c_ oscillations were less sensitive to negative regulation by PKC and independent of Ca^2+^ influx. Thus, differential feedback regulation by PKC gives rise to receptor-specific [Ca^2+^]_c_ oscillation profiles, which can encode the diverse physiological and pathophysiological responses to distinct agonists that all act through the IP_3_ signaling cascade.

## Introduction

Cytosolic Ca^2+^ ([Ca^2+^]_c_) oscillations are key regulators of cellular signaling and tissue physiology in a variety of cell types (1,2). In the liver, which is mainly comprised of hepatocytes, oscillatory [Ca^2+^]_c_ transients play a key role in bile secretion (3), regulation of mitochondrial oxidative phosphorylation (4), glucose metabolism (5,6), and tissue regeneration and gene expression (7). Oscillatory increases in [Ca^2+^]_c_ are well described in isolated hepatocytes and the intact liver challenged with hormones such as vasopressin and the catecholamines epinephrine and norepinephrine (8–12). These G-protein-coupled receptors (GPCRs) activate phosphoinositide-specific phospholipase C (PLC), resulting in the hydrolysis of phosphatidylinositol 4,5-bisphosphate (PIP_2_) into inositol 1,4,5-triphosphate (IP_3_) and diacylglycerol (DAG). IP_3_ mobilizes Ca^2+^ from the endoplasmic reticulum (ER) via IP_3_ receptors (IP_3_R), whereas DAG recruits and activates protein kinase C (PKC) to initiate specific protein phosphorylation cascades and a multitude of signaling outcomes (13,14). Modulation of PKC, by activation or inhibition of its isoforms, can modify hormone-induced [Ca^2+^]_c_ oscillation frequency and shape (15–17).

In hepatocytes, Ca^2+^ spike frequency is controlled by the agonist concentration, which encodes stimulus strength and determines the magnitude of downstream responses, whereas Ca^2+^ spike amplitude and kinetics are independent of the agonist dose (8,9,12,18,19). The [Ca^2+^]_c_ oscillation rising phase, driven by positive feedback of Ca^2+^ on the IP_3_R and PLC-β (20–22), is also relatively constant irrespective of the GPCR activated. However, there is agonist-specific diversity in the falling phase of the Ca^2+^ spikes, which gives rise to receptor-specific spike profiles that are remarkably distinguishable for each individual biological trigger (9,21,23). This Ca^2+^ spike decay phase sets the duration of the [Ca^2+^]_c_ transient, an important parameter that has been demonstrated to regulate activation of gene-specific transcription factors (24). Therefore, elucidation of the signaling machinery that regulates Ca^2+^ spike kinetics will further foster our understanding of physiological information encoded by [Ca^2+^]_c_ oscillations.

Extracellular nucleotides are key signaling molecules, recognized by hepatocytes and other liver cells types, affecting important hepatic processes (25). ATP binds to purinergic P2 receptors, a family of cell-surface receptors which have been divided into two classes based on their structures and modes of signal transduction; ligand-gated ion channels termed P2X receptors and G-protein-coupled receptors termed P2Y receptors. P2X receptors are ATP-gated ion channels permeable to Na^+^, K^+^ and Ca^2+^ cations (26). Seven subunits of these receptors (P2×1-7) expressed by different cell types are grouped either in a homomeric or heteromeric mode (27). P2Y receptors are metabotropic and activated by purines and pyrimidines, including ATP, ADP, UTP, UDP or UDP-glucose, and they are subclassified into P2Y1, 2, 4, 6, 11, 12, 13, and 14 subtypes (28) (P2Y11 receptor has been described only in humans (29)). P2Y1, 2, 4, 6, and 11 receptors are coupled to Gα_q_/_11_ G-proteins, leading to activation of the PLC - IP_3_ pathway and subsequent Ca^2+^ release from ER. P2Y12, 13 and 14 subtypes are coupled to G_i_ and G_o_, inhibiting adenylate cyclase and leading to decreased activity of cAMP-dependent protein kinases. Activation of these receptors does not result in a direct change in [Ca^2+^]_c_ (30).

Changes in purinergic signaling have been described in models of vascular injury, inflammation, insulin resistance, hepatic fibrosis, cirrhosis, diabetes, hepatitis, liver regeneration following injury or transplantation and cancer (31). Purinergic signaling is required for hepatocyte proliferation as knockout of the P2Y2 receptor impaired this process after partial hepatectomy (32). However, little is known about the physiological role of extracellular nucleotides in the healthy liver, and whether changes in purinergic signaling occur or contribute to liver pathophysiology.

The first reports of the characterization of extracellular nucleotide-induced [Ca^2+^]_c_ oscillations in hepatocytes were performed in primary rat cells microinjected with the Ca^2+^ indicator aequorin (23,33–37). These studies characterized the pharmacology of hepatic purinergic receptors involved in Ca^2+^ signaling and investigated downstream effects on metabolism. In the present study, we have used fura-2 Ca^2+^ imaging to revealed [Ca^2+^]_c_ oscillations similar to those previously described. Submaximal doses of endogenous purinergic agonists elicited discretely distinguishable [Ca^2+^]_c_ spike patterns for agonists known to act on different purinergic receptor subtypes. We found that the predominant P2Y receptors in rat hepatocytes were P2Y1, which is activated by ADP and yields short duration [Ca^2+^]_c_ spikes, and P2Y2 and P2Y4, which are activated by UTP to give longer duration more complex [Ca^2+^]_c_ spike patterns. ATP activates all three of these P2Y receptor subtypes and generates complex composite [Ca^2+^]_c_ oscillation patterns. An important question is what receptor-specific mechanisms give rise to the distinct stereotypic shape and duration of the individual [Ca^2+^]_c_ spikes? Previous studies have shown that rat hepatocytes also have functional P2X receptors that could contribute to the [Ca^2+^]_c_ oscillations in response to ATP (38). However, we found that the P2X receptors were not activated at the physiological doses of ATP that give rise to [Ca^2+^]_c_ oscillations in hepatocytes. Instead, our studies demonstrate that P2Y1 receptor activation with ADP elicits narrow [Ca^2+^]_c_ transients due to robust negative feedback by PKC, and that this effect of PKC is largely driven by Ca^2+^ influx at the plasma membrane. By contrast, the P2Y2/4 receptors activated by UTP generate broad [Ca^2+^]_c_ transients, which are less sensitive to PKC-dependent negative regulation and independent of Ca^2+^ influx.

## Results

### mRNA expression of P2 receptors in primary rat hepatocytes

cDNAs generated from five independent rat hepatocyte preparations were probed by RTqPCR using primers specific for P2X and P2Y receptors (Figure 1). Low mRNA expression levels of P2X receptors *P2×1*, *P2×2*, *P2×3*, *P2×5* and *P2×6* were detected whereas *P2×4* and *P2×7* transcripts were abundantly expressed in both freshly-isolated and overnight-cultured hepatocytes. For the G_αq_-coupled P2Y receptors, *P2y1*, *P2y2* and *P2y4* subtypes were abundantly observed, with very low transcript detection of *P2y6*. After overnight culture, a considerable decrease in *P2y4* expression was observed (Figure 1*B*), resulting in expression of predominantly *P2y1* and *P2y2* G_αq_-coupled receptors. The G_i_-coupled *P2y12*, *P2y13* and *P2y14* receptors were detected, albeit at a low expression levels at both time-points analyzed. Thus, according to the expression profile, mRNA transcripts mainly from *P2×4*, *P2×7*, *P2y1*, *P2y2* and *P2y4* were detected in rat hepatocytes, consistent with relevant physiological roles for these receptors.

**Figure 1.**
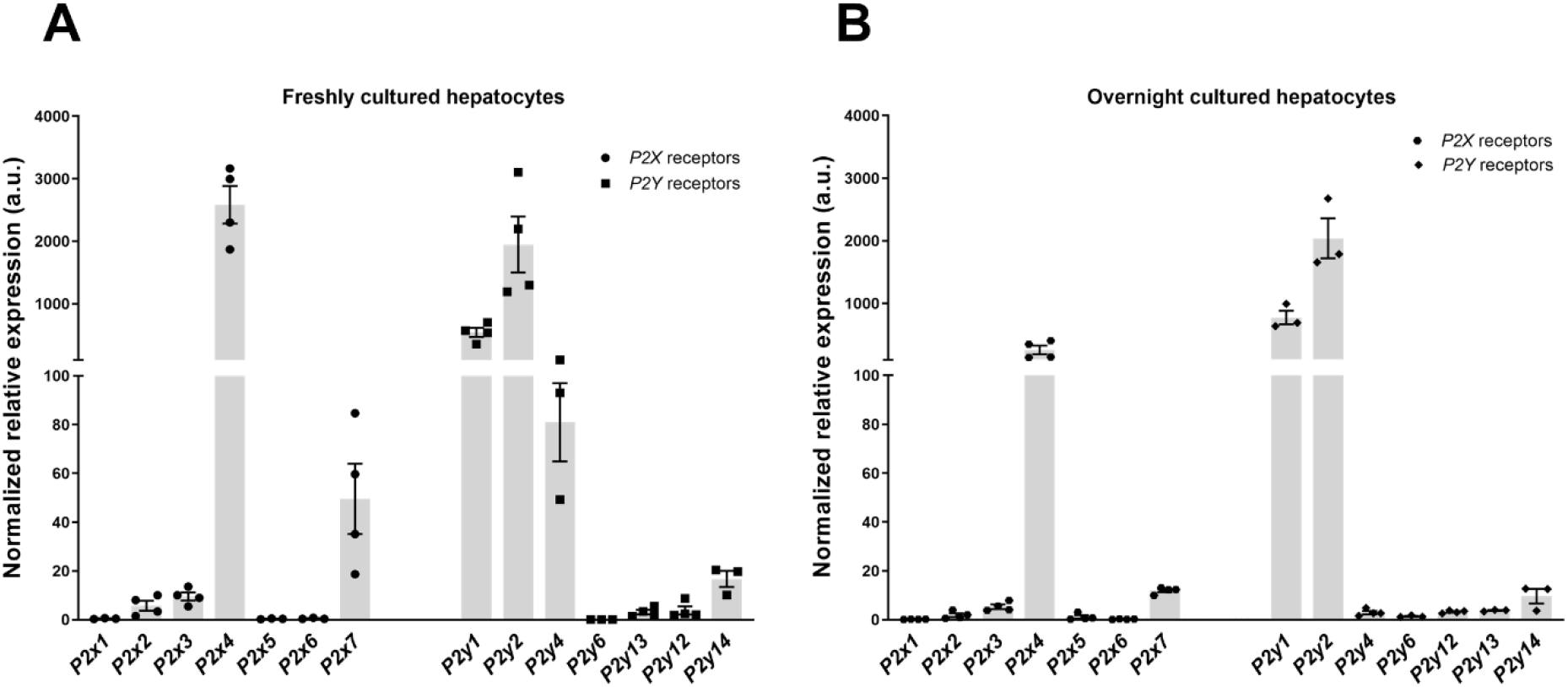
Purinergic P2 receptor gene expression in primary rat hepatocytes. RT-qPCR determination of mRNA expression levels of P2X and P2Y receptors from freshly isolated (**A**) and overnight (**B**) cultured hepatocytes. Purinergic receptor gene expression was normalized to *Rpl0* expression. Data are mean ± S.E.M from 3-4 hepatocyte preparations in each case.

### [Ca^2+^]_c_ oscillations characteristics and types of responses elicited by ATP, ADP and UTP

It has been previously reported that activation of purinergic receptors can generate a diverse pattern of [Ca^2+^]_c_ oscillations (23,36,37,39,40), although the specific contribution of each of the P2X and P2Y receptor types has not been fully described. In order to distinguish the Ca^2+^ responses elicited by different types of purinergic receptors, [Ca^2+^]_c_ oscillation responses to ATP, ADP, UTP and UDP were analyzed in primary rat hepatocytes. At low doses (1-5 μM) of physiological purinergic agonists, spike durations of the [Ca^2+^]_c_ oscillations were different, with distinguishable falling phases, indicating that the duration of the individual Ca^2+^ spikes is characteristic of the purinergic receptor being activated (Figure 2). Similar receptor-specific Ca^2+^ spike shapes have been reported for other G_αq_-coupled receptors, including V_1_-vassopressin and α_1_-adrenergic receptors (9,10,19). ATP, an agonist for all ionotropic P2X and metabotropic P2Y receptors, elicited complex Ca^2+^ spike shapes with two main patterns: broad spikes with small secondary oscillations resulting in a biphasic decay phase, and narrow spikes with a fast decay phase. Although both patterns can be observed in the same cell, broad Ca^2+^ spikes of relatively long duration were observed in the majority of ATP-stimulated hepatocytes. (Figure 2*A*). Spike width (measured as full width at half maximum; FWHM) of baseline-separated ATP-evoked Ca^2+^ spikes were 35.6 ± 3.5 s for the more complex broad spikes and 18.6 ± 2.1 s for the short duration spikes. ADP, an agonist of Gα_q_-coupled P2Y1 and G_i_-coupled P2Y12 and P2Y13 receptors, generated homogeneous short duration [Ca^2+^]_c_ oscillations (15.9 ± 0.9 s FWHM) with narrow peaks and a rapid declining phase, similar to the sporadic narrow Ca^2+^ spikes induced by ATP (Figure 2*B*). UTP, an agonist of P2Y2 and P2Y4 receptors, elicited predominantly longer duration [Ca^2+^]_c_ spikes (40.4 ± 3.5 s), similar to the broad long duration ATP responses (Figure 2*C*). UDP, a P2Y6 receptor-selective agonist, was the only extracellular nucleotide tested that failed to elicit a [Ca^2+^]_c_ response in rat hepatocytes (Figure 2*D*).

**Figure 2.**
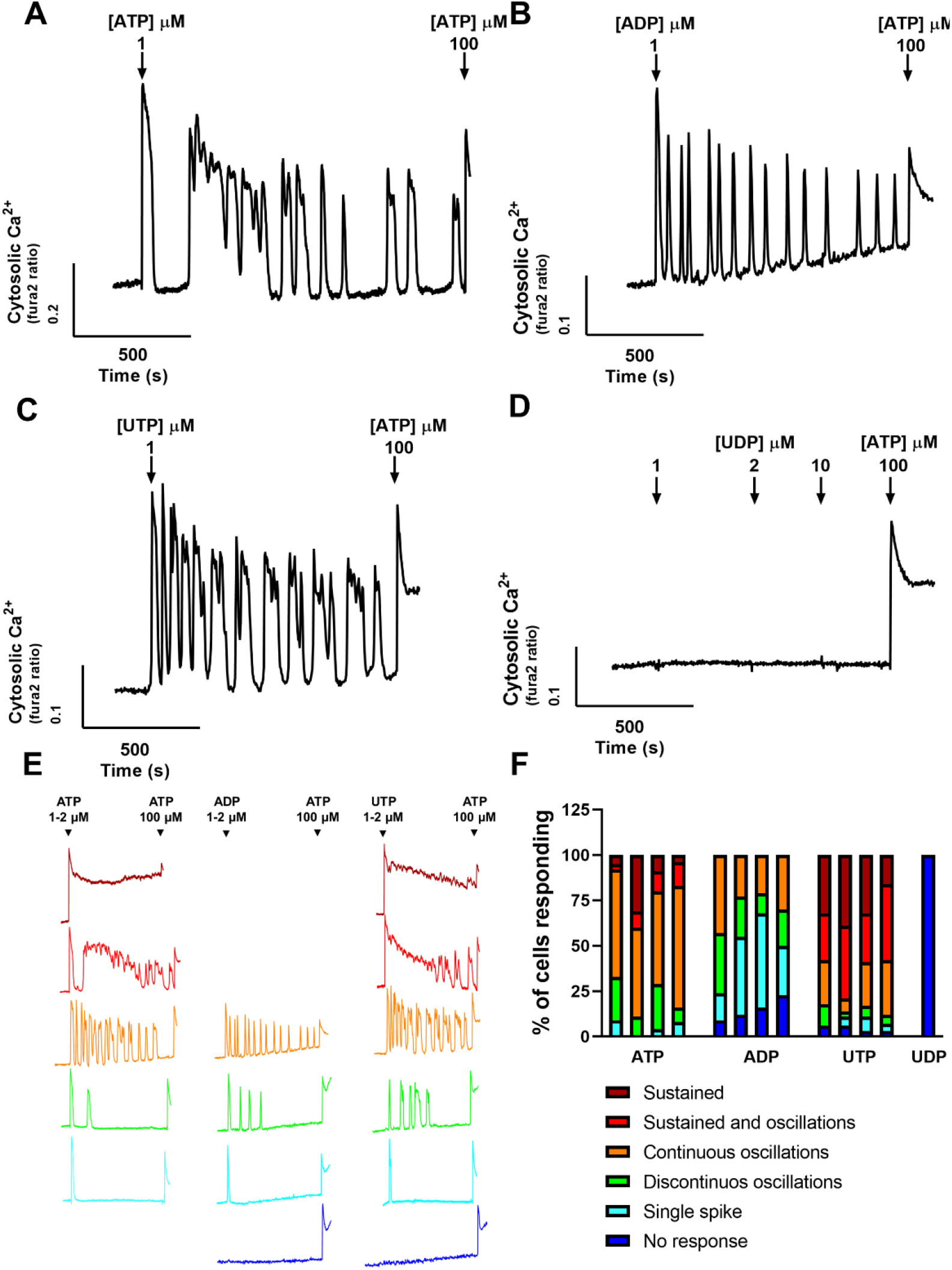
[Ca^2+^]_c_ oscillation profiles elicited by purine nucleotides in hepatocytes. Representative traces of typical ATP (**A**), ADP (**B**), UTP (**C**), and UDP-induced (**D**) [Ca^2+^]_c_ responses in hepatocytes loaded with fura-2. The indicated concentration of each nucleotide was added at the arrows and remained present for the remainder of the experiment. **E:** At low agonist doses (1-2 μM), different strengths of Ca^2+^ response from *No response* (blue), *Single spike* (cyan), *Discontinuous oscillations* (green), *Continuous Oscillations* (orange), *Sustained & oscillations* (red) through to *Sustained* (dark red) can be elicited by each extracellular nucleotide. A maximal dose of ATP (100 μM) was added at the end of each trace. **F:** Ordinal plot of the Ca^2+^ response strength in hepatocytes challenged with ATP, ADP, UTP or UDP. (Data are from ≥ 50 cells in each of four independent experiments).

The signal strength for Ca^2+^-dependent hormones in liver is not a function of [Ca^2+^]_c_ amplitude, which is relatively constant, but is encoded in the frequency, duration and robustness of the [Ca^2+^]_c_ oscillations (8,9,21). These [Ca^2+^]_c_ responses can be classified using an ordinal scale ranging from no response, through single spikes, then oscillations and up to a sustained [Ca^2+^]_c_ increase. Stimulation of different receptors by each extracellular nucleotide evoked not only distinct Ca^2+^ spike profiles, but also a different range in the magnitude of [Ca^2+^]_c_ response (Figures 2, *E* and *F*). ATP was able to elicit all types of [Ca^2+^]_c_ responses, from single spikes to a fully sustained increase (Figure 2*E*, left panel), and was the only nucleotide able to evoke an increase in [Ca^2+^]_c_ in all analyzed cells (Figure 2*F*). P2Y2 and P2Y4 receptor activation by UTP generated mostly large initial peaks followed by a sustained plateau and repetitive broad [Ca^2+^]_c_ spikes (Figures 2, *E*, right panel, and *F*). At the same agonist dose, activation of P2Y1 receptors by ADP evoked only a range of single, irregular or repetitive narrow [Ca^2+^]_c_ transients without any sustained [Ca^2+^]_c_ signals (Figures 2, *E*, middle panel, and *F*).

### P2X receptors do not contribute to [Ca^2+^]_c_ oscillations in rat hepatocytes

The similarity of the oscillatory [Ca^2+^]_c_ responses induced by ATP and UTP to those elicited by other Gα_q_-coupled receptors (8,9) suggests that these purinergic Ca^2+^ signals arise from IP_3_-dependent mobilization of intracellular Ca^2+^ stores through activation of P2Y receptors. However, since abundant mRNA expression of the P2×4 and P2×7 receptors was observed, we designed experiments to determine whether P2X receptors contribute to the observed [Ca^2+^]_c_ signals. We used YM-254890, a specific inhibitor for Gα_q_ (41) to block P2Y receptor coupling to this G-protein. Addition of 1 μM YM-254890 reversed the [Ca^2+^]_c_ response elicited by 5 μM ATP (Figure 3 *A*), and blocked all [Ca^2+^]_c_ responses to ATP concentrations of 1-100 μM (Figure 3, A and *B*). At higher ATP concentrations (300-400 μM), a slow [Ca^2+^]_c_ increase to a plateau level was observed (Figures 3, *B* and *C*), suggesting that P2X receptor activation and Ca^2+^ influx through the associated plasma membrane channels only occurs at very high ATP concentrations. ATP never elicited [Ca^2+^]_c_ oscillations in the presence of YM-254890. Consistent with a role for P2X receptors at high ATP concentrations, the P2×4/P2×7 receptor agonist BzATP (10-20 μM) also caused a monophasic [Ca^2+^]_c_ increase in the presence of YM-254890 (Figures 3, *B* and *C*), and similar to ATP, BzATP did not cause [Ca^2+^]_c_ oscillations under these conditions. Even when added in the absence of YM-254890, 10 μM BzATP caused a sustained [Ca^2+^]_c_ increase without oscillations (Figure 3*D*). At lower concentrations of BzATP [Ca^2+^]_c_ oscillations were sometimes observed in the absence of YM-254890, consistent with the partial agonist activity of BzATP against P2Y1 receptors (38,42). Taken together, these observations indicate that the ATP-induced [Ca^2+^]_c_ oscillations in rat hepatocytes are evoked by P2Y receptors, specifically P2Y1 and P2Y2/4, through the IP_3_-dependent intracellular Ca^2+^ signaling pathway, and do not depend on P2X receptors.

**Figure 3.**
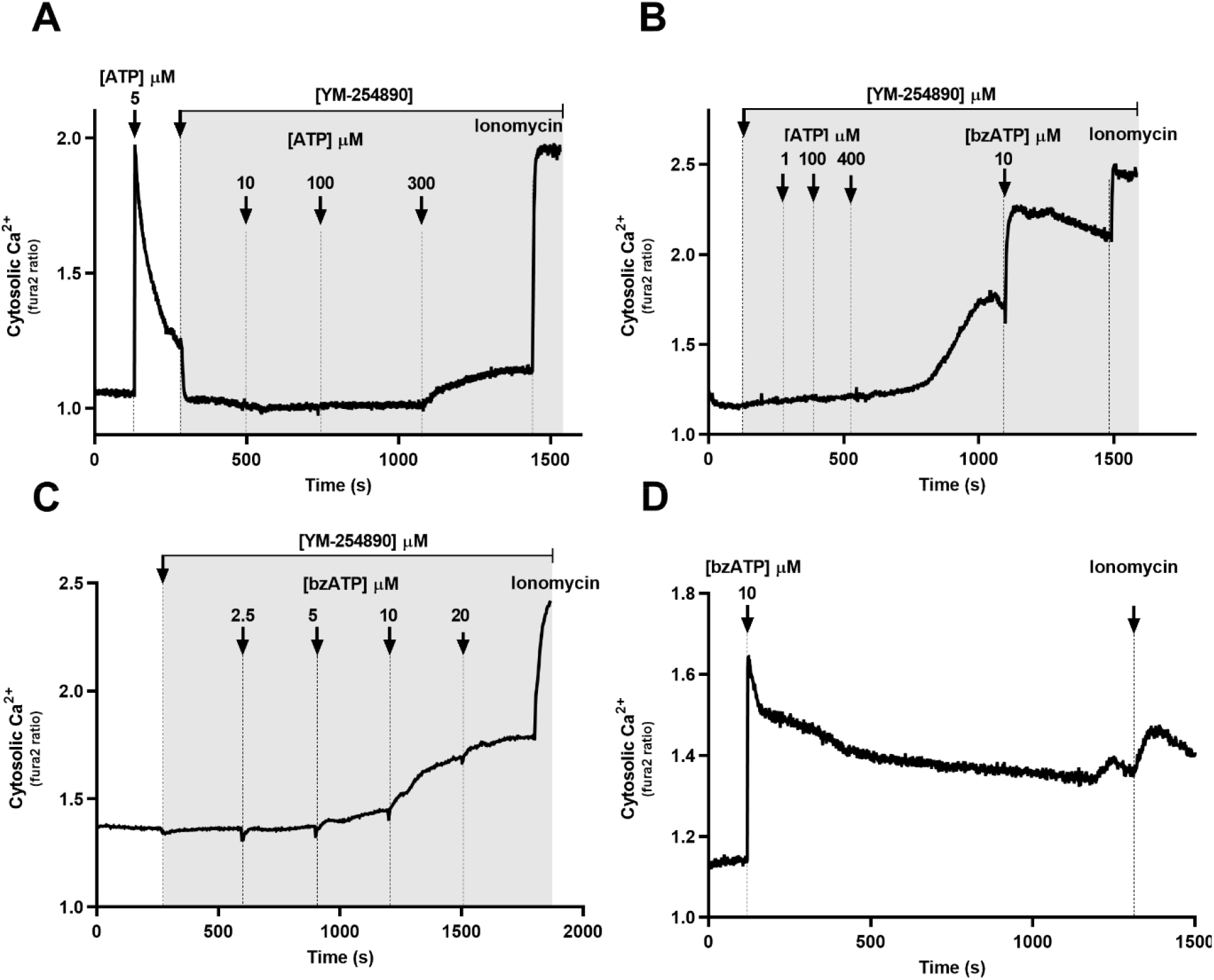
P2X receptor activity does not contribute to [Ca^2+^]_c_ oscillations in hepatocytes. Hepatocytes loaded with fura-2 were exposed to the indicated concentrations of ATP (in μM) added at the arrows and remaining present throughout each experiment. The Gα_q_ protein inhibitor YM-254890 (1 μM) was present during the period indicated by the grey shading. (**A**) ATP (5 μM) induced a large [Ca^2+^]_c_ spike that was terminated by addition of YM-254890. Subsequent additions of increasing concentrations of ATP (10 and 100 μM) had no effect on [Ca^2+^]_c_, whereas a high ATP dose (300 μM) caused a slow monophasic [Ca^2+^]_c_ increase. (**B**) Effect of increasing ATP concentrations in the presence of YM-254890, followed by addition of the P2X agonist BzATP (10 μM). **(C)** Dose response to BzATP in the presence of YM-254890. (**D**) Response to BzATP (10 μM) in the absence of YM-254890.

### Downregulation of PKC increases the Ca^2+^ spike width for ADP- and UTP-induced [Ca^2+^]_c_ oscillations

The [Ca^2+^]_c_ oscillations elicited by activation of P2Y1 and P2Y2/4 receptors have characteristically distinct Ca^2+^ spike shapes, which could suggest distinguishable physiological roles in the liver. The molecular mechanisms that contribute to distinct P2Y receptor-induced [Ca^2+^]_c_ oscillation shapes are still poorly understood. It has been shown previously that PKC activation or inhibition can have multiple effects on hormone-induced Ca^2+^ oscillation kinetics and frequency in hepatocytes (15–17). We examined the effect of downregulation of conventional and novel PKC isoforms by overnight treatment (16-24 h) with 1 μM phorbol 12-myristate 13-acetate (PMA), referred to as PKC-DR, or the inactive analogue 4α-PMA, referred to as CTR. In both ADP- and UTP-stimulated PKC-DR cells, the oscillation frequency was reduced and the spike width (FWHM) prolonged when compared to control cells (Figure 4). Although PKC-DR showed similar effects for both purinergic receptor agonists, the magnitude of the effect was not the same. PKC-DR cells stimulated with ADP showed a 1.8-fold increase in spike duration (17.5 ± 0.5 s to 31.1 ± 0.9 s; CTR and PKC-DR, respectively; *p* < 0.0001) (Figure 4*C*), whereas UTP stimulation of PKC-DR cells resulted in a more modest 1.3-fold increase in spike width (27.7 ± 0.5 s to 35.5 ± 0.7 s; CTR and PKC-DR, respectively; *p* < 0.001) (Figure 4*H*). Notably, the basal spike width for ADP in control (CTR) cells is about 10 s shorter than for UTP (*p* <0.0001), but the spike widths and overall [Ca^2+^]_c_ oscillation shapes became more similar after PKC-DR (Figure 4). PKC-DR also had a differential effect on the magnitude of the [Ca^2+^]_c_ response elicited by P2Y1 and P2Y2/4 receptors in hepatocytes. Specifically, PKC-DR resulted in a more robust ADP response profile, with a shift to fewer unresponsive cells and an increase in oscillatory and sustained [Ca^2+^]_c_ responses (Figure 4*E*). By contrast, there was no shift in the pattern of [Ca^2+^]_c_ response signatures in PKC-DR cells after UTP challenge (Figure 4*J*). The differential susceptibility of P2Y receptor-dependent [Ca^2+^]_c_ oscillations to PKC downregulation could indicate that P2Y1 receptors are more sensitive to negative feedback inhibition by PKC than P2Y2/4 receptors (see Discussion).

**Figure 4:**
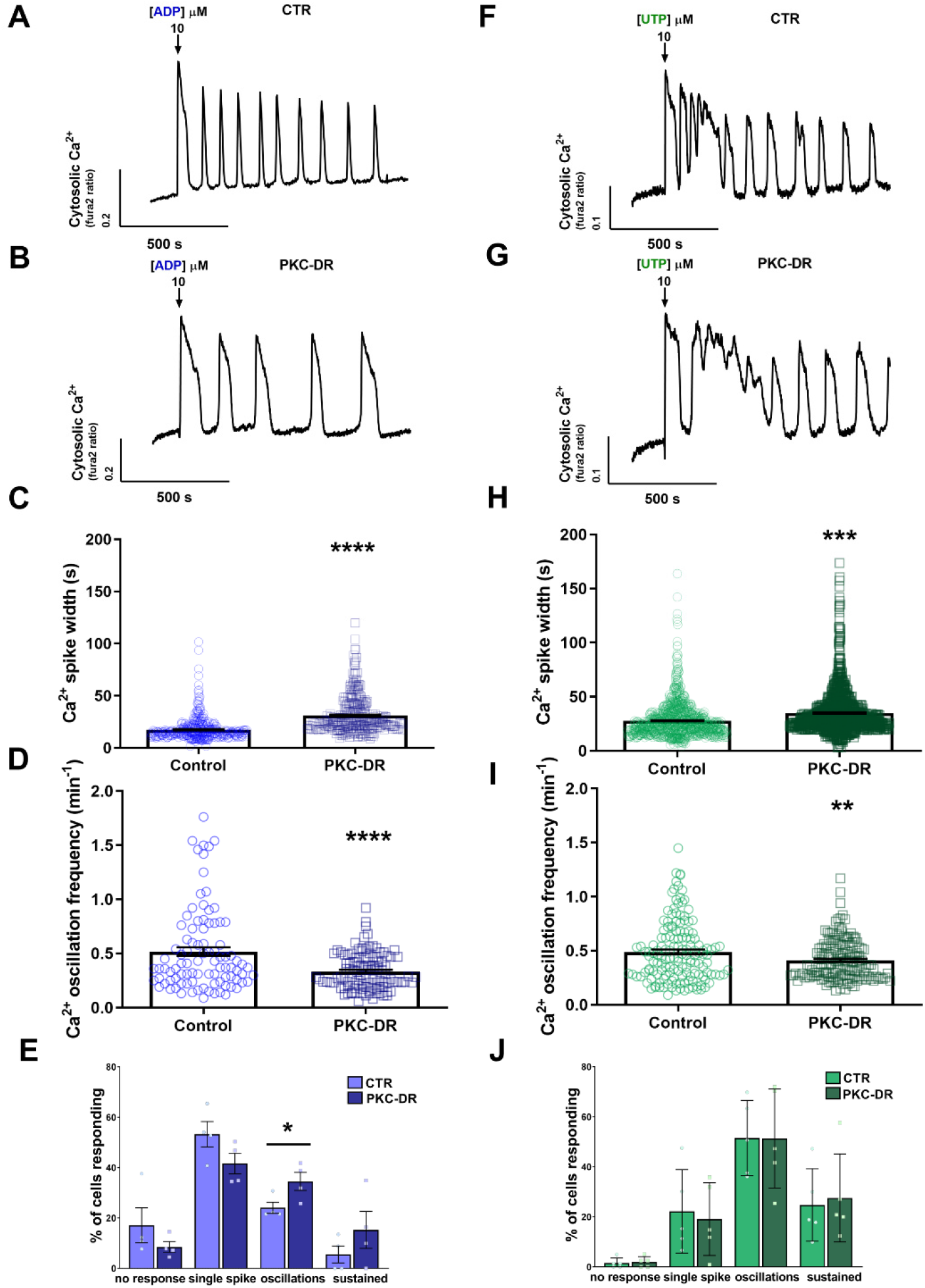
Downregulation of PKC prolongs ADP- and UTP-induced Ca^2+^ spike duration in isolated rat hepatocytes. Isolated hepatocytes were cultured overnight with PMA (1 μM) to downregulate conventional and novel PKC isoforms (***PKC-DR***) or with the inactive analogue α-PMA (1 μM, ***CTR***). The cells were loaded with fura-2 and then stimulated with ADP or UTP (10-15 μM). Representative traces for ADP- and UTP-induced [Ca^2+^]_c_ responses are shown for control (**A-F**) and PKC-DR hepatocytes **(B-G)**. Summary data show the effects of PKC-DR on Ca^2+^ spike width measured as full width at half maximum (FWHM) and oscillation frequency for ADP (**C-D**) and UTP (**H-I**) induced [Ca^2+^]_c_ transients. The distribution of Ca^2+^ response patterns are also shown for ADP (**E**) and UTP (**J**). Blue and green symbols represent data from ADP- and UTP-induced [Ca^2+^]_c_ responses, respectively. Data are mean ± S.E.M. from ≥ 50 cells from at least three independent experiments. *, *p*<0.05; **, *p*<0.01; ***, *p*<0.001; **** *p*<0.0001; Student’s *t* test.

### Extracellular Ca^2+^ is required for negative feedback inhibition of [Ca^2+^]_c_ oscillations by PKC

To determine the impact of plasma membrane Ca^2+^ influx on P2Y-evoked [Ca^2+^]_c_ oscillations, we compared Ca^2+^ spike kinetics in the presence and absence of extracellular Ca^2+^. Purinergic agonist-induced [Ca^2+^]_c_ oscillations were monitored in hepatocytes incubated in medium containing the normal physiological 1.3 mM CaCl_2_ or switched to Ca^2+^ free buffer just prior to data acquisition (representative traces are shown in Figures 5, *A-B* and *D-E*). In the absence of extracellular Ca^2+^, the individual ADP-evoked Ca^2+^ spike widths were longer (35.64 ± 1.16 s) than those observed in the presence of extracellular Ca^2+^ (18.84 ± 1.09 s) at the same agonist dose (Figure 5, *A*–*C*). In UTP-stimulated cells, no difference was observed in Ca^2+^ spike width in the presence (29.25 ± 0.79 s) or absence of extracellular Ca^2+^ (30.79 ± 1.41 s) (Figure 5*F*). Although P2Y2/4 receptor activation with UTP in Ca^2+^-free conditions did not result in a changed of the Ca^2+^ spike width, the shape of the falling phase was qualitatively different. The characteristic UTP-induced Ca^2+^ spikes with a biphasic decay phase containing secondary small Ca^2+^ spikes (Figure 5*D*) (see also Figure 2*C*) were not observed in the absence of extracellular Ca^2+^, resulting in a less complex falling phase (Figure 5*E*).

**Figure 5:**
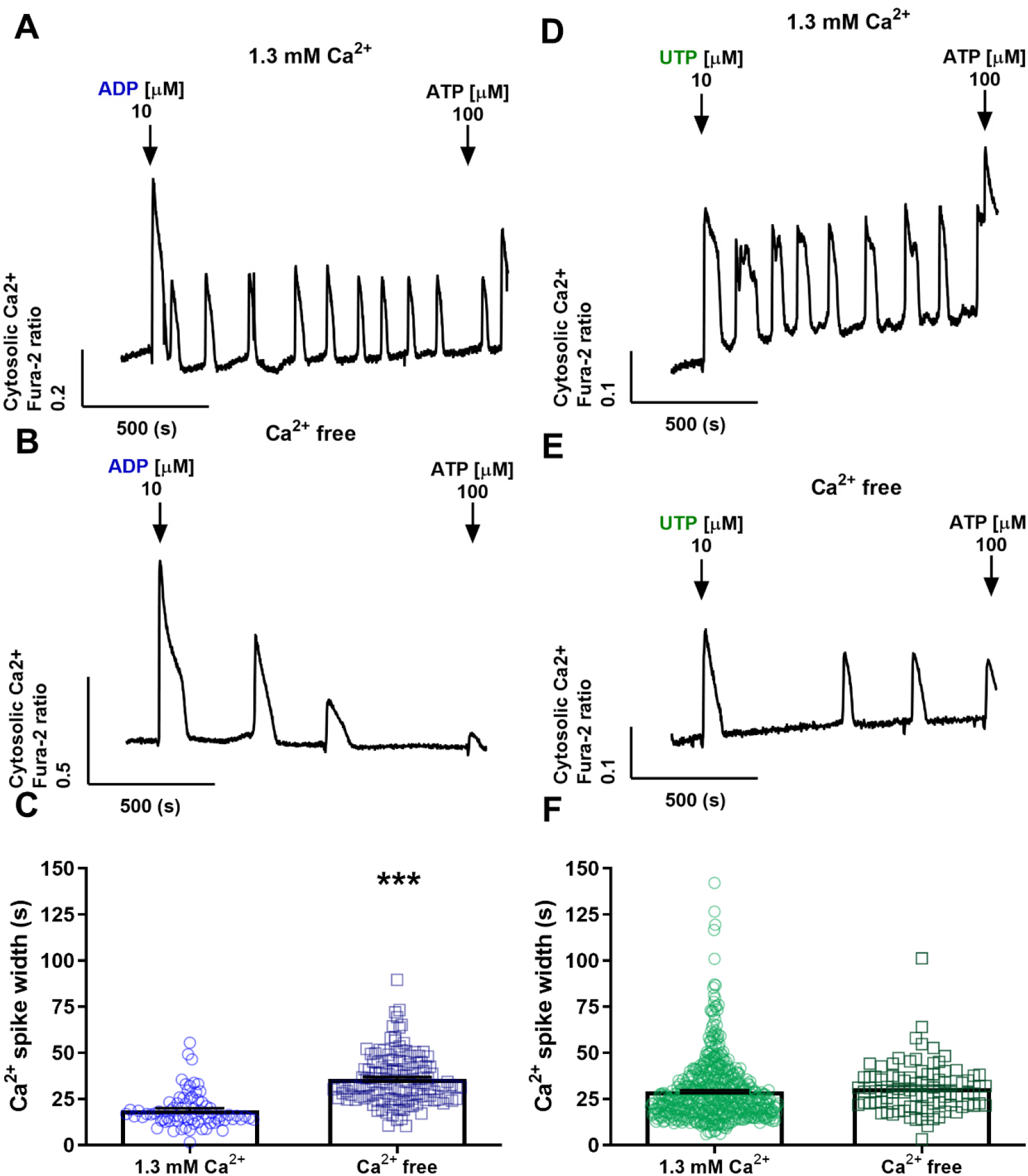
Absence of extracellular Ca^2+^ differentially affects P2Y1 and P2Y2/4 receptor-dependent [Ca^2+^]_c_ oscillations. Isolated hepatocytes were loaded with fura-2 and then stimulated with ADP and UTP (1-10 μM) in the presence (***1.3 mM Ca*^*2*+^**) or absence (***Ca*^*2*+^ *free***) of extracellular Ca^2+^. Representative traces of typical ADP (**A-B)** and UTP (**D-E)** evoked [Ca^2+^]_c_ responses are shown. Summary data of the effect of extracellular Ca^2+^ removal on Ca^2+^ spike width (FWHM) for ADP (**C**) and UTP (**F**). Blue and green symbols represent data from ADP- and UTP-induced Ca^2+^ spikes, respectively. Data are mean ± S.E.M. from ≥ 50 cells from at least three independent experiments. ***, *p*<0.001; Student’s *t* test.

The data described above suggest that plasma membrane Ca^2+^ entry is required for negative feedback inhibition of P2Y1-dependent [Ca^2+^]_c_ oscillations elicited by ADP, but not for the P2Y2/4-dependent [Ca^2+^]_c_ oscillations elicited by UTP. Thus, for P2Y1 receptors, ADP stimulation after PKC-DR or in the absence of extracellular Ca^2+^ resulted in longer duration [Ca^2+^]_c_ spikes, whereas P2Y2/4 receptor-dependent responses were affected to a lesser extent following PKC-DR and were unaffected by removal of extracellular Ca^2+^. One possible explanation for the selective effect of extracellular Ca^2+^ on the P2Y1 responses is that plasma membrane Ca^2+^ entry could be required for negative feedback inhibition of P2Y1 receptors by PKC. To determine whether the absence of extracellular Ca^2+^ disturbs PKC-dependent regulation of Ca^2+^ spike kinetics elicited by the different P2Y receptor types, we investigated whether PKC-DR affects ADP- and UTP-induced [Ca^2+^]_c_ responses in the presence or absence of extracellular Ca^2+^ (representative traces are shown in Figures 6, *A-B* and *D-E*). In PKC-DR cells, [Ca^2+^]_c_ spikes elicited by P2Y1 or P2Y2/4 receptor activation were found to have the same duration with or without extracellular Ca^2+^ (Figures 6, *C* and *F*). These data showing that extracellular Ca^2+^ does not affect Ca^2+^ spike width when PKC activity is downregulated, suggest that Ca^2+^ entry at the plasma membrane is important for the PKC-dependent regulation of oscillatory [Ca^2+^]_c_ signals.

**Figure 6:**
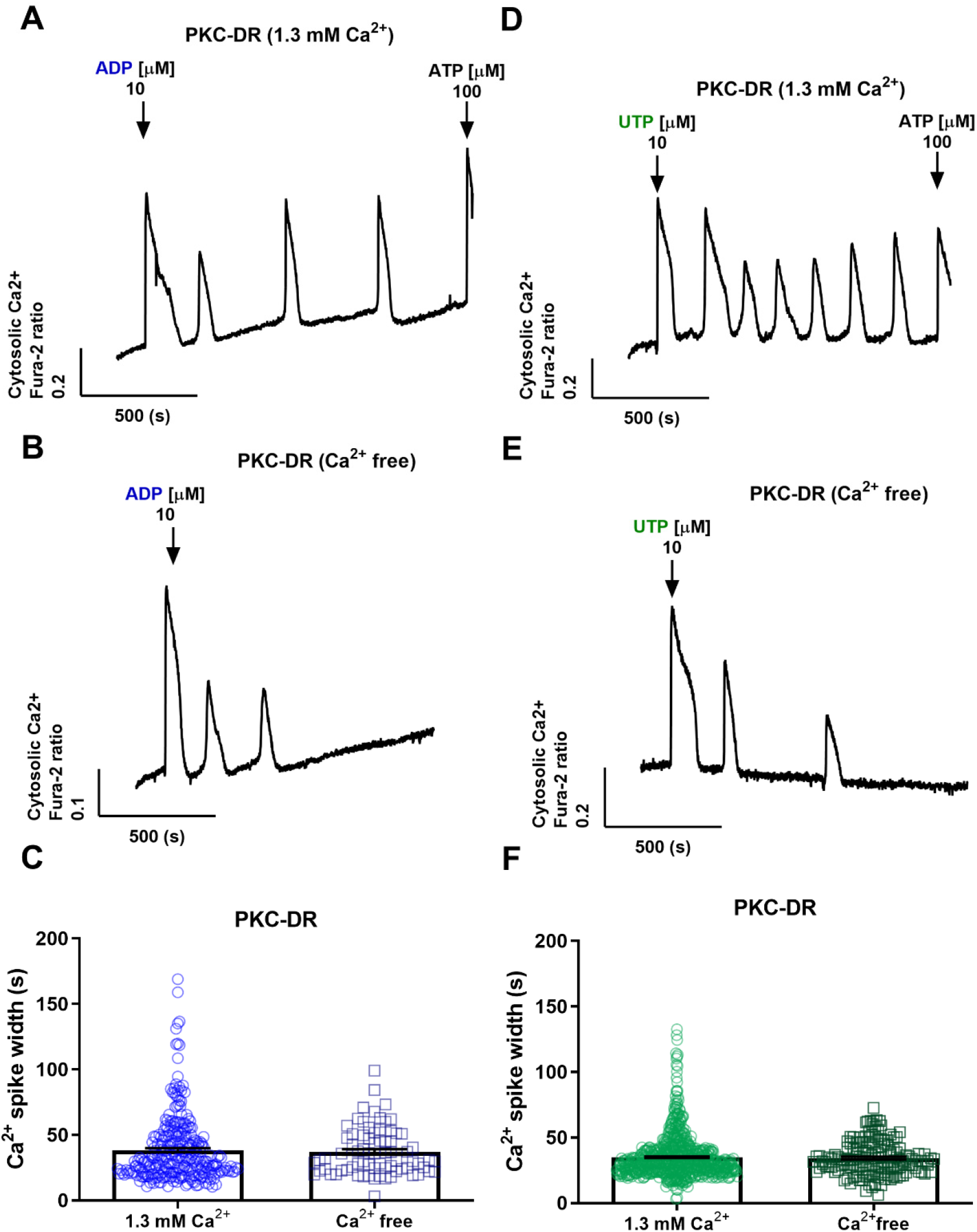
Extracellular Ca^2+^ has no effect on [Ca^2+^]_c_ oscillation spike width following PKC downregulation. Isolated hepatocytes were cultured overnight with PMA (1 μM) to downregulate conventional and novel PKC isoforms (PKC-DR). The cells were then loaded with fura-2 and stimulated with ADP or UTP (10-15 μM) in the presence (***1.3 mM Ca*^*2*+^**) or absence (***Ca*^*2*+^ *free***) of extracellular Ca^2+^. Representative traces for ADP (**A-B**) and UTP (**D-E**) are shown under both experimental conditions. Summary data of the effect of extracellular Ca^2+^ removal on Ca^2+^ spike width (FWHM) in PKC-DR hepatocytes is shown for ADP (**C**) and UTP (**F**). Blue and green symbols represent data from ADP- and UTP-induced [Ca^2+^]_c_ oscillations, respectively. Data are mean ± S.E.M. from ≥ 50 cells from at least three independent experiments; Student’s *t* test.

### Acute effect of PKC activation and inhibition on ADP and UTP-induced [Ca^2+^]_c_ transients

To further investigate the role of PKC in the regulation of hepatic purinergic receptor Ca^2+^ signaling, we examined the effect of acute activation and inhibition of PKC on ADP- and UTP-induced [Ca^2+^]_c_ oscillations. Hepatocytes were stimulated with low doses of ADP or UTP (1-5 μM), to elicit [Ca^2+^]_c_ oscillations and then the effect of PKC modulators was determined in the same cells (representative traces are shown in Figures 7, *A-B* and *G-H*). [Ca^2+^]_c_ oscillation frequency and spike width (FWHM) were calculated in cells that displayed continuous [Ca^2+^]_c_ oscillations for at least 5 minutes before and after application of the PKC modulators. For P2Y1 receptors stimulated with ADP, activation of PKC by PMA decreased [Ca^2+^]_c_ oscillation frequency (Figure 7*C*), with no change in the spike width (Figure 7*D*). By contrast, for P2Y2/4 receptors stimulated with UTP, PKC activation with PMA did not change the [Ca^2+^]_c_ oscillation frequency (Figure 7*I*), but caused a decrease in spike width (Figure 7*J*). Acute inhibition of PKC with bisindolylmaleimide (BIM) had no effect on ADP- or UTP-induced [Ca^2+^]_c_ oscillation frequency (Figures 7, *E* and *L*). However, BIM elicited opposite effects on the width of the Ca^2+^ spikes evoked by ADP and UTP. The spike width for ADP-induced [Ca^2+^]_c_ oscillations increased from 16.33 ± 1.05 s to 32.88 ± 1.55 s (*p* < 0.0001) (Figure 7*F*), whereas spike width for UTP-induced [Ca^2+^]_c_ oscillation decreased from 36.36 ± 0.65 s to 20.45± 0.29 s (*p* < 0.0001) (Figure 7*M*). A similar effect on Ca^2+^ spike duration was observed with staurosporine, a nonselective inhibitor of protein kinases, including protein kinase C (43). Treatment with 500 nM staurosporine caused a small increase in the ADP-induced Ca^2+^ spike width (from 15.12 ± 0.94 s to 17.95 ± 0.77 s, *p* < 0.001), and a more pronounced decrease in Ca^2+^ spike width with UTP (from 56.24 ± 2.98 s to 22.21 ± 0.66 s, *p* < 0.0001) (data not shown). Thus, although off-target effects of BIM and staurosporine are possible, the broadening of ADP-induced Ca^2+^ spikes and narrowing of UTP-induced Ca^2+^ spikes with both of these PKC inhibitors are consistent. Taken together, the data with acute and chronic manipulation of PKC demonstrate that this is an important feedback pathway that plays a key role in shaping the [Ca^2+^]_c_ oscillations elicited by purinergic agonists, and that it does so in a receptor-specific manner.

**Figure 7:**
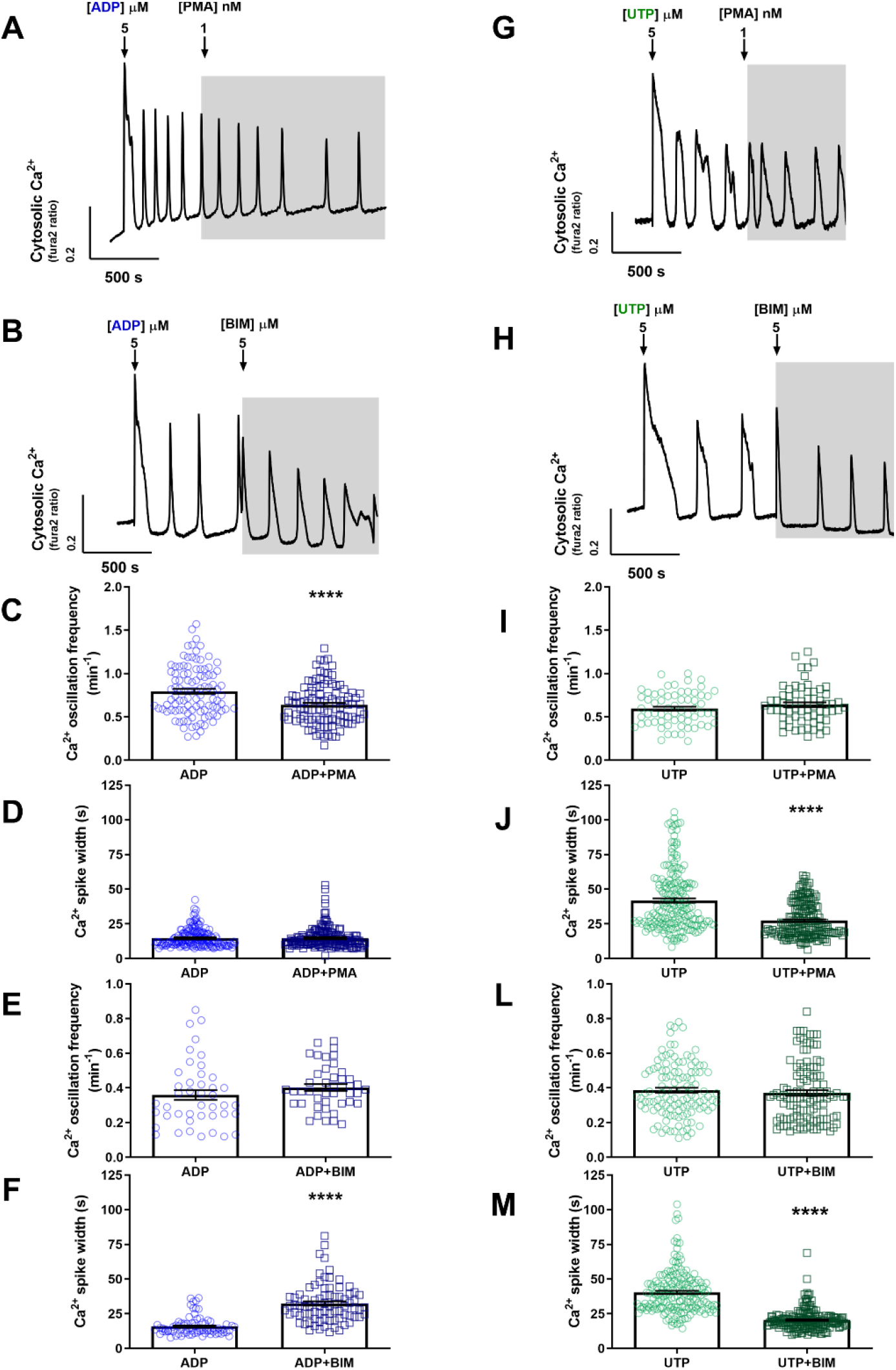
Effects of acute activation and inhibition of PKC on ADP- and UTP-induced [Ca^2+^]_c_ oscillations. The effects of PMA (1 nM) and BIM (5 μM) on ADP- and UTP-induced [Ca^2+^]_c_ oscillations were analyzed in hepatocytes loaded with fura-2. After stimulation with the purinergic agonists, cells were treated with PMA or BIM as indicated by the shaded areas on the Ca^2+^ traces. Representative [Ca^2+^]_c_ oscillation responses are shown for ADP (**A-B)** and UTP (**G-H**). The frequency and width of the Ca^2+^ spikes induced by ADP and UTP were calculated from 5-min periods in the absence or presence of the PKC modulators. Summary data are shown for oscillation frequency and spike width (FWHM) for ADP (**C-F)** and UTP (**I-M**). Blue and green symbols represent data from ADP- and UTP-induced [Ca^2+^]_c_ oscillations, respectively. Data are mean ±S.E.M. from ≥ 50 cells from at least three independent experiments. ****, *p*<0.0001; Student’s *t* test.

## Discussion

The physiological actions of extracellular nucleotides in the liver include regulation of bile secretion, glucose homeostasis and cell regeneration. At the cellular level, ATP can be released by hepatocytes into basolateral, sinusoidal or apical extracellular compartments, acting as potent autocrine and paracrine signals to regulate liver physiology (25,44). ATP is also released as a cotransmitter with norepinephrine from sympathetic nerves innervating the liver, and serves to stimulate glycogenolysis and gluconeogenesis through a Ca^2+^-dependent signaling pathway (25). Under pathophysiological conditions, local release of nucleotides and/or circulating levels of ATP can increase, disturbing the balance of purinergic signaling in the liver (45). In the present study we defined the P2X and P2Y receptors expressed in rat hepatocytes and analyzed [Ca^2+^]_c_ oscillations evoked by the predominant P2Y1 and P2Y2/4 receptors. The physiological signals encoded by these complex [Ca^2+^]_c_ dynamics were shown to be regulated by distinct PKC feedback mechanisms on the P2Y receptors, without any contribution of P2X receptors.

As demonstrated here and by others (38,46), P2×4 and P2×7 are the most abundantly expressed P2X receptor isoforms in rat hepatocytes. Immunohistochemistry of intact rat liver showed P2×4 receptor localized in both the basolateral and apical canalicular domains of the hepatocyte (46), consistent with the reported role of P2×4 receptors in biliary secretion (47). Previous studies have used BzATP, an agonist of P2×4 and P2×7 receptors, to show that these receptors are functional in hepatocytes, including the activation of a Na^+^-dependent inward current and Ca^2+^ influx in a rat hepatoma cell line (38,46), and increased [Ca^2+^]_c_ and enhanced small molecule permeability in isolated rat hepatocytes (38,46). In addition, the P2×4 receptor allosteric activator ivermectin also increases [Ca^2+^]_c_ in hepatoma cells (38,46).

In primary rat hepatocytes, low doses of ATP (1-5 μM), the agonist for all P2 receptors, elicits oscillatory [Ca^2+^]_c_ transients (23,37,48). Gonzalez and colleagues demonstrated previously that treatment with high ATP (1 mM) or BzATP (100-300 μM) induced membrane pore formation and blebbing in rat hepatocytes, and these events were inhibited by oxidized ATP, an antagonist of P2×7 receptors (38). In the present study, we demonstrated that [Ca^2+^]_c_ responses elicited by low (1-10 μM) concentrations of ATP are blocked by a Gα_q_ inhibitor, whereas the sustained monophasic [Ca^2+^]_c_ increases observed at high ATP concentrations are unaffected by Gα_q_ inhibition. It seems likely that P2×7 receptor activity accounts for these monophasic [Ca^2+^]_c_ responses, since activation of this receptor has been shown to require higher ATP concentrations (EC_50_ ≥ 100 μM) compared to other P2X receptors (49,50). However, since BzATP activates both P2×4 and P2×7 receptors (51), the pharmacological approach cannot distinguish between the individual roles of these receptors on the sustained [Ca^2+^]_c_ increase in rat hepatocytes at high levels of ATP.

Summarizing the findings with respect to P2X receptors, although P2×4 and/or P2×7 are functional and able to increase [Ca^2+^]_c_ in rat hepatocytes, these receptors do not appear to play a role in the generation of [Ca^2+^]_c_ oscillations and, therefore, in the physiological liver functions mediated by these Ca^2+^ signals. Nevertheless, their activation by high ATP levels suggests a potential role of P2X purinergic signaling under pathophysiological conditions in liver. Indeed, a role of the P2×7 receptor in cytotoxicity and ATP mediated apoptosis has been described in the liver (38,52). Importantly, perfusion of rat liver with ATP, within the dose range shown in this study to induce oscillatory [Ca^2+^]_c_ responses (1-10 μM ATP), increases hepatic glucose output, indicating that P2Y receptors are responsible for stimulating glycogenolysis (53).

In the present study, P2Y1 and P2Y2 receptor mRNA were the most abundantly expressed among the Gα_q_ coupled P2Y receptors, in agreement with previous mRNA expression analysis from rat hepatocytes (37). Although mRNA expression of P2Y6 receptors has been reported from non-quantitative PCR analysis (37), our quantitative PCR data revealed only a low expression level of P2Y6 receptors compared to other P2Y subtypes. At the functional level, UDP, which is the most potent P2Y6 agonist (54), failed to elicit a [Ca^2+^]_c_ response, consistent with previous findings (37). All of the other endogenous purinergic nucleotide agonists, ATP, ADP, and UTP, were able to elicit oscillatory [Ca^2+^]_c_ increases. Rat P2Y2 and P2Y4 receptors can be activated by ATP and UTP equipotently (55,56), so the relative roles of P2Y2 and P2Y4 receptors cannot be distinguished based on the specificity of UTP action. However, since we report 25-fold greater P2Y2 mRNA expression level than P2Y4 in freshly isolated cells, it is likely that the [Ca^2+^]_c_ oscillations elicited by UTP are mediated predominately by P2Y2 receptors. In our studies, ATP was the only agonist able to evoke a response in 100% of hepatocytes. The inability of ADP and UTP to elicit a [Ca^2+^]_c_ response in all cells could be explained by different expression levels of P2Y1 and P2Y2/4 receptors, which could relate to heterogeneous distribution of these receptors through the liver, particularly along the zonal axis of the hepatic lobule.

The oscillatory [Ca^2+^]_c_ signaling induced by Gα_q_-coupled receptors is well established to regulate hepatic metabolism (4–6). Furthermore, individual GPCRs elicit [Ca^2+^]_c_ oscillations with distinct Ca^2+^ spike kinetics, most notably in the falling phase, allowing for differential regulation of downstream targets (9,19,37). Several components of the Ca^2+^ signaling pathway have been described that modulate [Ca^2+^]_c_ oscillations and the profile of individual Ca^2+^ spikes via positive and negative feedback mechanisms. Positive feedback of [Ca^2+^]_c_ on PLC and consequent cross-coupling of Ca^2+^ and IP_3_ is an essential component in the generation of [Ca^2+^]_c_ oscillations in hepatocytes (20–22). In addition, negative feedback by PKC on PLC-dependent IP_3_ formation plays a role in spike termination, and in setting the frequency of hormone-induced [Ca^2+^]_c_ oscillations (15). The remarkably distinct Ca^2+^ spike profiles evoked by P2Y1 and P2Y2/4 receptors suggests that differential regulation of the Ca^2+^ mobilization machinery gives rise to receptor-specific differences in the duration and kinetics of the Ca^2+^ spike falling phase.

The narrow Ca^2+^ spikes associated with P2Y1 receptor activation and the broad Ca^2+^ spikes associated with P2Y2/4 receptor activation are both altered by PKC modulation, but with clear differences that contribute to the distinct Ca^2+^ spike profiles. Inhibition of PKC activity by PKC-DR increased the spike width in all cases, but this was much more pronounced for P2Y1 receptor. This PKC-DR approach has also been shown to enhance the oscillatory [Ca^2+^]_c_ responses to the α-adrenergic agonist phenylephrine in rat hepatocytes, again with a prolongation of the Ca^2+^ spike width (15). In those studies we demonstrated that agonist-stimulated PLC activity and IP_3_ production were enhanced due to the absence of negative feedback by PKC on the GPCR-dependent stimulation of PLC (15). Thus, the broadening of [Ca^2+^]_c_ oscillations observed for activation of P2Y1 and P2Y2/4 receptors are also likely due to the suppression of PKC negative feedback on IP_3_ production, particularly during the falling phase of the individual Ca^2+^ spikes. We interpret this to suggest that under normal conditions there is a larger element of PKC negative feedback onto the P2Y1 as opposed to the P2Y2/4 receptors, such that PKC-DR has a much more pronounced effect to modulate the P2Y1 response to ADP. Consistent with this, PKC-DR shifted the [Ca^2+^]_c_ signature profile toward more oscillatory and sustained responses for ADP stimulation, but not for UTP stimulation (Figures 4E vs 4F). This indicates that there is a selective effect of PKC to suppress the strength of the [Ca^2+^]_c_ signals elicited by P2Y1 receptor activation, which is relieved by elimination of this negative feedback after PKC-DR.

The differential PKC-dependent feedback mechanism of P2Y1 and P2Y2/4 receptors was also evidenced by acute modulation of PKC. P2Y1 activation with ADP in the presence of the PKC inhibitor BIM led to broader [Ca^2+^]_c_ spike widths, consistent with the data obtained with PKC-DR. As discussed above, this can be explained by the elimination of PKC negative feedback on the P2Y1 receptor-stimulated PLC. By contrast, acute PKC activation with PMA did not further decrease the already narrow Ca^2+^ spikes with P2Y1 receptor activation, but did slightly decrease the [Ca^2+^]_c_ oscillation frequency. These data are consistent with a strong endogenous negative feedback effect of PKC on P2Y1 receptors. As noted above, there was only a small effect of PKC-DR on P2Y2/4-induced [Ca^2+^]_c_ oscillations, suggesting that endogenous PKC-dependent negative feedback plays a lesser role than for the P2Y1 responses. Nevertheless, acute PKC activation with PMA caused a significant narrowing of UTP-induced [Ca^2+^]_c_ oscillations, demonstrating that P2Y2/4 receptor-mediated PLC activation is susceptible to PKC negative feedback but this is not full engaged by the endogenous activation of PKC during UTP stimulation. The paradoxical finding that BIM also caused narrowing of the Ca^2+^ spike width in response to UTP stimulation could be explained by a discrete site of action, perhaps through a different PKC isozyme that is not susceptible to PKC-DR. One potential target is the IP_3_R, which is sensitized to IP_3_ by PKC activation but is unaffected by PKC-DR in hepatocytes (15). Thus, with P2Y1 receptor activation the predominant effect of PKC may be negative feedback on PLC-dependent IP_3_ generation at the plasma membrane, whereas with P2Y2/4 receptor activation intracellular IP_3_R sensitization by PKC may predominate.

Our findings with respect to the influence of extracellular Ca^2+^ on the shape of the [Ca^2+^]_c_ oscillations elicited by P2Y1 and P2Y2/4 receptor activation sheds some light on the differential effects of PKC. Bearing in mind the importance of plasma membrane Ca^2+^ entry in maintaining [Ca^2+^]_c_ oscillations, the observation that the Ca^2+^ spikes elicited by ADP were substantially broader in the absence of extracellular Ca^2+^ was unexpected. Moreover, this effect was specific to P2Y1 receptor activation by ADP, and was not observed with UTP activation of P2Y2/4 receptors. The effect of extracellular Ca^2+^ to broaden the [Ca^2+^]_c_ spike width appears to be mediated by PKC, since there was no additional effect of Ca^2+^ removal in hepatocytes after PKC-DR. In other cell types there is evidence that extracellular Ca^2+^ is important for PKC activation, and Ca^2+^ influx may play a direct role in the translocation and activation of plasma membrane-associated conventional PKC isoforms (57–59). Our observation in hepatocytes that plasma membrane Ca^2+^ entry is required for negative PKC feedback only on ADP-induced [Ca^2+^]_c_ oscillations is consistent with the conclusion that PKC acts selectively on P2Y1 receptor signaling to shape the narrow Ca^2+^ spikes elicited by this receptor, whereas P2Y2/4 receptors signaling is refractory to this feedback. It is also significant because it provides evidence that plasma membrane Ca^2+^ entry can have discrete effects on PKC activation, distinct from the concomitant changes in bulk cytosolic Ca^2+^.

From a broader perspective, these data indicate that multiple PKC isoforms with distinct Ca^2+^ signaling targets and different modes of activation are engaged to fine tune agonist-induced [Ca^2+^]_c_ oscillations and to shape the Ca^2+^ spike kinetics for a given GPCR. Indeed, in human platelets, distinct PKC isoforms have been shown to regulate P2Y1 and P2Y12 receptor function and trafficking. Overexpression of dominant-negative PKC-α and PKC-δ isoforms revealed both novel and conventional PKC-mediate P2Y1 desensitization, whereas only novel PKCs regulate P2Y12 receptors (60). Another possible PKC target in the control of IP_3_ formation and Ca^2+^ mobilization is PLC. PKC-mediated phosphorylation of PLC-β3 (but not PLC-β1) in response to P2Y2 and M3 muscarinic receptor activation has been shown to decrease PLC association with Gα_q_/_11_ and contribute to the termination of the [Ca^2+^]_c_ increases evoked by these receptors (61). This type of negative feedback on PLC activity could explain the effects of PKC on Ca^2+^ spike duration observed in the present work. PKC activity can also regulate the degradation of IP_3_. In fibroblast cell lines, PKC phosphorylates IP_3_ Kinase isoforms A and B, decreasing the Ca^2+^/calmodulin-stimulated activity (62), thereby slowing the removal of IP_3_. However, this would not explain our findings with P2Y1 receptors, where PKC activity is associated with shorter duration Ca^2+^ spikes. By contrast, in platelets PKC activates IP_3_ 5-phosphatase, increasing the rate of IP_3_ degradation (63) and inhibition with staurosporin increases IP_3_ accumulation (64). Thus, the prolonged Ca^2+^ spikes seen with P2Y1 receptor activation after PKC-DR or acute PKC inhibition could be explained by lack of IP_3_ 5-phosphatase activation, although it is more difficult to see how this would result in a differential effect on P2Y1 vs P2Y2/4 receptors. Finally, the IP_3_R Ca^2+^ channel itself appears to be a PKC target. We have shown that the frequency of [Ca^2+^]_c_ oscillations elicited by uncaging IP_3_ in hepatocytes is increase by acute PMA treatment, indicating that IP_3_-induced Ca^2+^ release may also be enhanced by PKC (15). Overall, these multiple actions of PKC, and perhaps others, contribute to regulate the frequency and individual spike dynamics of agonist-induced [Ca^2+^]_c_ oscillations.

Taken together, our results show that P2Y1 and P2Y2/4 receptors are functional and display stereotypic [Ca^2+^]_c_ oscillations that are tightly regulated by PKC feedback mechanisms in primary rat hepatocytes. P2Y1 receptor-evoked [Ca^2+^]_c_ oscillations seem to be shaped by a strong negative feedback on PLC activation, with a key role of plasma membrane Ca^2+^ entry in this component of PKC action. The distinct [Ca^2+^]_c_ oscillation pattern seen with P2Y2/4 receptor activation appears to involve differential regulation by PKC, reflecting differences in the sensitivity of IP_3_ generation and downstream components of IP_3_ metabolism and action in the Ca^2+^ signaling cascade. The P2Y receptor-specific [Ca^2+^]_c_ signatures regulated by PKC in the liver provide a means to regulate the diverse downstream targets, including both physiological and pathophysiological processes, all encoded by the complex [Ca^2+^]_c_ oscillation signals.

## Experimental procedures

### Primary Cell Culture

Isolated hepatocytes were prepared by collagenase perfusion of livers obtained from male Sprague-Dawley rats. Cells were maintained in Williams E medium for 1–6 h for experiments using freshly isolated cells or 16–24 h for experiments using overnight cultured cells, as described previously (9,18). Animal studies were approved by the Institutional Animal Care and Use Committee at Rutgers, New Jersey Medical School.

### RNA extraction and cDNA synthesis

Total RNA was isolated from hepatocytes using TRIzol reagent and followed by column purification (Qiagen), according to the manufacturer protocol. DNAse I treatment (Ampgrade, 1U/μg of RNA, Thermo Fisher Scientific) was performed for 15 min at room temperature to prevent residual DNA contamination. RNA was quantified by spectrophotometry (NanoDrop, Thermo Fisher Scientific,). Two micrograms of DNAse-treated RNA of each sample were simultaneously reverse transcribed using Superscript™ III First-Strand Synthesis System (Thermo Fisher Scientific) according to the manufacturer protocol. After cDNA synthesis, samples were submitted to a 20-minute digestion with RNAseH at 37° C.

### Quantitative PCR

Quantitative transcript analyses were performed in a StepOnePlus™ Real-Time PCR System (Thermo Fisher Scientific), as described previously (65). Optimal conditions were obtained using a five-point, two-fold cDNA and primer dilution curve for each amplicom. Each qPCR reaction contained 12.5 ng of reversely transcribed RNA, each specific primer at 200 nM (Table 1) and SYBR Green PCR Master Mix (Thermo Fisher Scientific), following the manufacturer conditions. Samples with no DNA or with RNA (no reverse transcription) were included as negative controls. A dissociation curve was acquired to confirm product specificity and the absence of primer dimers. Relative transcript amount quantification was calculated from three technical replicates, as previously described (66,67). Purinergic receptor gene expression was normalized to *Rpl0* expression, which did not change under the used experimental conditions.

**Table 1:**
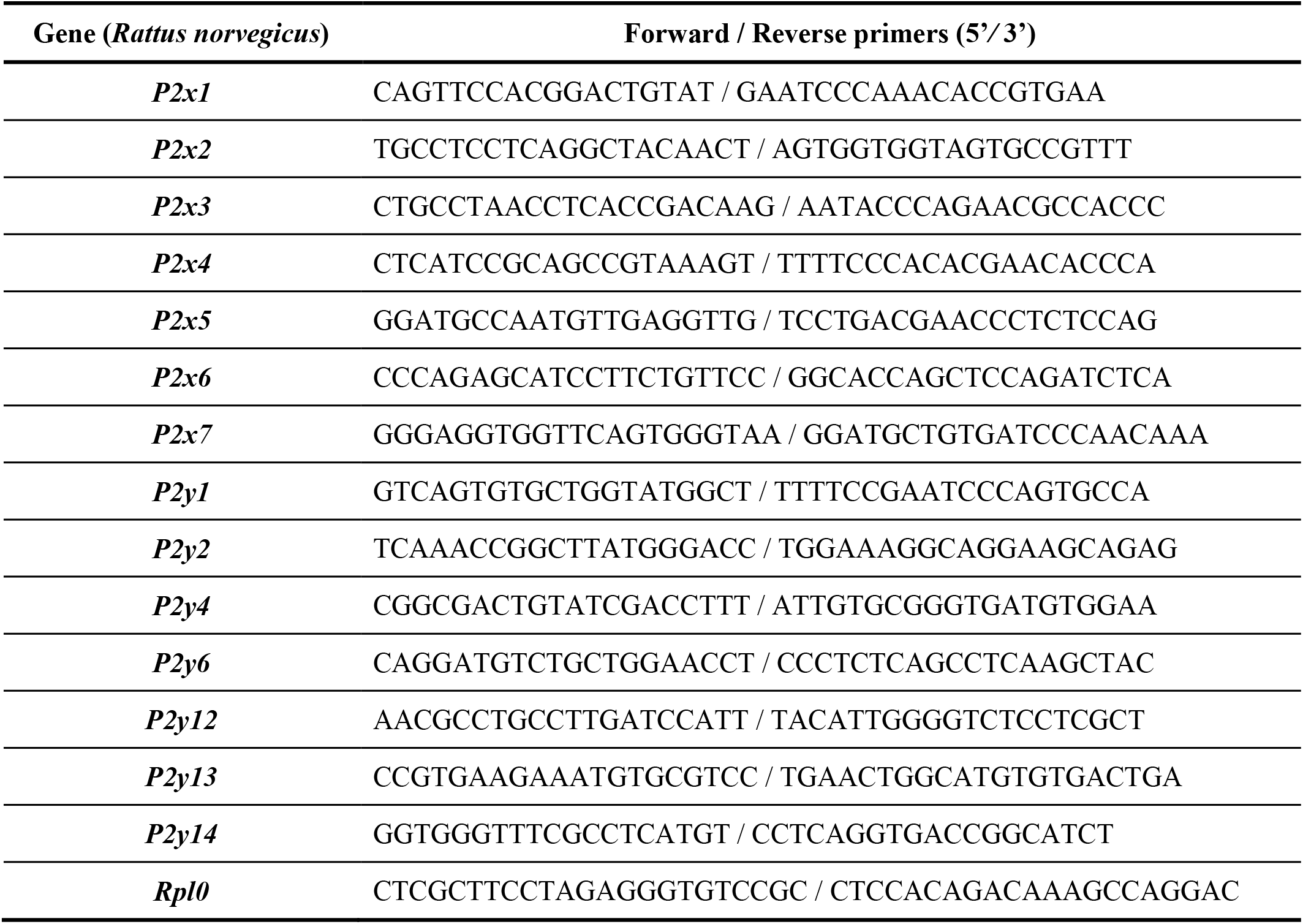
Gene-specific primers used for RTqPCR.

### Cytosolic Ca^2+^ Measurements

Calcium imaging experiments were performed in HEPES-buffered physiological saline solution comprised of 25 mM HEPES (pH 7.4), 121 mM NaCl, 5 mM NaHCO_3_, 10 mM glucose, 4.7 mM KCl, 1.2 mM KH_2_PO_4_, 1.2 mM MgSO_4_, 1.3 mM CaCl_2_, and 0.25%(w/v) fatty acid-free BSA and supplemented with the organic anion transport inhibitors sulfobromophthalein (100 μM) or probenecid (200 μM) to increase retention of fura-2. Hepatocytes were loaded with fura-2 by incubation with 5 μM fura-2/AM and Pluronic acid F-127 (0.02% v/v) for 20–40 min. Cells were transferred to a thermostatically-regulated microscope chamber (37° C). Fura-2 fluorescence images (excitation, 340 and 380 nm, emission 510 nm long pass) were acquired at 1 to 3 s intervals with a cooled charge-coupled device camera coupled to an epifluorescent microscope, as described previously (68).

### Data Analysis

Relative expression of purinergic receptors genes was calculated according to (66). Briefly, the arithmetic means of replicated cycling threshold (Cq) value of each gene was transformed to a quantity taking into account the amplification efficiency of each gene. The raw quantities were subsequently normalized to the reference gene. For the imaging data, the frequency and spike width (full width at half maximum, FWHM), were determined using algorithms (Brumer R *et al*, unpublished work) written in MATLAB (MathWorks, Natick, MA, USA). Graph plotting and data analysis were performed with GraphPad Prism software. Statistical analysis was performed using two-tailed Student’s *t* test.

## Data availability

All data are contained within the manuscript.

## Funding and additional information

Supported by the Thomas P. Infusino Endowment and NIH R01DK078019. HU acknowledges grant support from the São Paulo Research Foundation (FAPESP project 2018/07366-4) and fellowship support from CNPq (CNPq project 306392/2017-8).

